# A modulator of cognitive function: Cerebellum modifies human cortical network dynamics via integration

**DOI:** 10.1101/2024.09.12.612716

**Authors:** Kanika Bansal, Zaira Cattaneo, Viola Oldrati, Chiara Ferrari, Emily D. Grossman, Javier O. Garcia

## Abstract

The cerebellum, with distinctive architecture and extensive cortical connections, has long been associated with motor control; however, evidence suggests its role extends beyond motor functions, playing a crucial role in cognitive processes. Despite these insights, how cerebellar computations modulate cortical networks remains elusive. Here, we evaluate dynamic network reconfigurations in the cerebral cortex connectivity following noninvasive inhibitory repetitive transcranial magnetic stimulation (rTMS) targeting the right cerebellum. Using dynamic community detection, we uncover the dynamic network properties by which cerebellar stimulation spreads through the cortex, inspecting the evolution of modular network structures prior to and after cerebellar stimulation. Our results indicate that: (1) *flexibility*, or the likelihood of network nodes to change module allegiances, increases post stimulation; (2) dynamic patterns in which module allegiances emerge and evolve are individualistic and do not follow a single functional prototype; and (3) cerebellar nodes play the role of integrators for distinct network modules. These results suggest that the cerebellum plays a pivotal role in modulating distributed cortical activity, seamlessly integrating and segregating information beyond motor control. This integrative capacity may underlie the cerebellum’s contributions to high-level cognitive functions and, more broadly, to the foundation of human intelligence.

## Introduction

The neural underpinnings of cognition – and more generally *intelligence* – are rooted in the intricate interplay of several complex cognitive and neural systems. Initially understood to solely support highly skilled motor control, the cerebellum is now recognized as modulating cognitive, language, social and affective functions (Leiner et al., 1993; Schmahmann, 2010; Buckner, 2013; Schmahmann et al., 2019; Cattaneo et al., 2022). Cognitive impairments observed after cerebellar lesions - termed *dysmetria of thought -* manifest in the regulation of speed, capacity, and appropriateness of cognitive processes, especially in forming predictions that may improve the efficiency with which cognitive processes are executed (Honda et al., 2018; Van Overwalle et al., 2020). This disruption parallels the cerebellum’s role in mediating highly skilled motor behaviors through the fine-tuning of movement execution (Koziol et al., 2014). The means by which computational mechanisms in the cerebellum promote efficiency in cortical processing are not yet fully understood.

Cerebellar cytoachitectonic organization is strikingly unique, with a highly uniform and tightly folded, or foliated, structure that contains many more neurons than the cortical sheet (Andersen et al., 1992; Herculano-Houzel, 2010). Cerebellar Purkinje cells possess significantly more dendritic spines than typical cortical cells, allowing for the broad integration of inputs at a speed and scale beyond the operating parameters of the cerebral cortex, providing a computational vehicle to rapidly integrate cortical information (Walter and and Khodakhah, 2009). It is widely understood that the way we learn fine motor movements within our complex and dynamic world is by way of the creation and use of an internal model of effectors operating within the environment, in large part supported by cerebellar mechanisms (Honda et al., 2018). Much in the same way, the cerebellum has been linked to predicting and modulating cognitive processes, optimizing performance as a function of the current environment, particularly in the context of tasks that emphasize timing demands (Koziol et al., 2014).

Each microfold of the cerebellum is organized as a closed loop circuit with a single cortical target, the fundamental unit of cerebro-cerebellar architecture that is the basis of the universal cerebellar transform theory (Leiner et al., 1993; Schmahmann and Pandya, 1997; Strick et al., 2009). This theory posits that the many functional subdivisions of the cerebellum execute the same computation, namely adaptive control via integration of internal models with incoming sensory cues, and it is the cortical target of the cortico-cerebellar loop that determines any functional specialization (Leiner et al., 1986; Schmahmann et al., 2019). Consistent with this proposal, cortical expansion of the cerebellum over evolution tracks with that of cerebral cortex (Smaers, 2014), with a notable exception of over-representation of frontoparietal control networks in the cerebellum relative to cerebral cortex (Marek et al., 2018). More recent proposals emphasize the potential for the cerebellum to coordinate long-range corticocortical coherence by aligning the relative phase of local ongoing neural oscillations between distal cortical regions (Georgescu Margarint et al., 2020; McAfee et al., 2022). These findings demonstrate how local activity in the cerebellum has the potential to co-modulate long-range functionally connected targets in the cerebral cortex.

Neurostimulation approaches, such as transcranial magnetic stimulation (TMS) and transcranial electrical stimulation (tES), have proven instrumental for revealingthe roles and dynamics of cortico-cortical and cerebellar-cortical interactions. These methods enable precise modulation of the underlying local neural activity along with distal, functionally connected circuits, uncovering causal relationships with cognitive systems. For instance, studies like Rastogi et al. (2017) demonstrated that inhibitory theta burst stimulation over Crus I, a cognitive processing hub in the lateral portion of the cerebellum (Marek et al., 2018), selectively reduces connectivity with the prefrontal cortex’s frontal control regions but has no impact on neural activity in the hand region of motor cortex. Likewise, excitatory neuromodulation delivered to Crus 1 of the lateral cerebellum increases connectivity to the cortical default mode network (DMN), distinct from midline vermis stimulation, which increased connectivity to the dorsal attention network (DAN), highlighting the function-specific propagation of cerebellar neuromodulation to cortical circuits (Esterman et al., 2017; Halko et al., 2014).

Complementing neurostimulation, advancements in network neuroscience, such as dynamic community detection, allow for the characterization of both local and global neural changes especially when paired with neurostimulation techniques (e.g., (Garcia et al., 2020a, 2020b, 2018)), offering insights into how neural communities reorganize in response to stimulation. This suite of tools and those belonging to the more general *network neuroscience*, have demonstrated the cerebellum to have small-world properties, showing that the cerebellum – similar to the cerebral cortex – balances regular connectedness with random connectivity, and consequently, facilitating nuanced interplay between neural circuits (Chen et al., 2022). Together, these approaches provide an integrated strategy by which the intricate interactions that underpin brain functionality may be revealed, and may be fruitful in understanding the cerebellum’s foundational functional principles of motor control but also the influence on cognitive processes, more generally.

In this study, we employed a “perturb and measure” approach using noninvasive cerebellar stimulation to explore its effects on cerebral cortex dynamics. By administering 1 Hz inhibitory rTMS to the right cerebellum Crus I region and subsequently analyzing resting state fMRI data in individuals, we were able to compare network structures and activity before and after stimulation. Specifically, our analysis focused on changes in brain network dynamics due to cerebellar stimulation, including shifts in the overarching community structure and variations in nodal properties focusing on how integration, recruitment, and the roles of dynamic hubs and integrators within these networks are affected by targeted cerebellar manipulation.

## Methods

### Experimental data acquisition

Nineteen participants that met the inclusion criteria for participation in both the TMS and MRI procedures were recruited for this experiment. Two participants were excluded from the analysis because they did not participate in all experimental sessions. Hence, seventeen individuals (3 males; M = 25.6 SD = 1.9 years old) were included in the final sample. All subjects provided informed consent as approved by the local Institutional Review Board.

#### Procedure

Participants completed three experimental sessions on three separate days. Each session for each participant was scheduled at the same time of the day; each consecutive session was interleaved by a minimum of 2 days to a maximum of 15 days. In session 1 participants participated in single-pulse stimulation to assess their resting state motor threshold. In session 2, participants underwent two consecutive resting-state scans, during which T1 and T2*- weighted images were acquired. In session 3, participants completed two resting state scans, one pre and one post the offline inhibitory rTMS protocol.

#### Stimulation

In session 1, TMS was administered using a MagProx100 stimulator (MagVenture) connected to a butterfly-shaped coil, with static cooling (MCF-B70 coil, MagVenture). Single-pulse TMS was applied over the left primary motor cortex (M1) at increasing intensities to determine each participant’s resting motor threshold (rMT). The rMT was defined as the minimum stimulation intensity required to elicit motor-evoked potentials (MEPs) with an amplitude of at least 50 mV in the first dorsal interosseous muscle with a 50% probability (Rossini et al., 1994). In session 3, participants received 1 Hz rTMS for 20 minutes over the right cerebellum Crus I (Talairach coordinate 22, −75, −21) at 100% resting motor threshold as assessed in session 1 and in line with previous TMS studies targeting the cerebellum (Demirtas-Tatlidede et al., 2011; Ferrari et al., 2018). The cerebellar target region was identified by neuro-navigated TMS (using the neuronavigation system SofTaxic, EMS srl, Italy) on each participant’s individual magnetic resonance image acquired in session 2. Offline inhibitory stimulation was delivered using a refrigerated coil connected to the stimulator, and the rTMS coil was positioned tangentially to the scalp, parallel to the midsagittal line, with the handle pointing superiorly, following evidence that this orientation effectively modulates cerebellar activity (Bijsterbosch et al., 2012; van Dun et al., 2017).

#### Image acquisition

MR images were acquired using a 3T (Philips Healthcare, Best, The Netherlands) scanner with a 32-channel head coil.In session 2, T1-weighted images were collected for each individual [192 images, TR = 10 sec, TE = 101 ms]. Subjects then participated in a single 18:40 min resting state scan with the instructions to lay quietly with eyes open during which T2*-weighted images were collected [560 volumes, TR = 2 sec, TE = 27 ms]. In session 3, participants repeated the resting state scan two additional times, once prior to and once immediately following the rTMS stimulation. Post-stimulation fMRI image acquisition was initiated within M = 2.8, SD = 0.49 min (range 2.5 −4 min) after cessation of the stimulation.

#### Data preprocessing

All functional images were preprocessing in BrainVoyager (Brain Innovations, Inc), including corrected for head motion within the scan, spatial smoothing (4mm FWHM gaussian kernel), linear detrending and temporal high-pass filtering (cutoff frequency: 0.01 Hz). Functional images were then registered to the individual subject volumetric T1-weighted anatomical images. Cortical regions of interest were identified using the Schaefer template (Schaefer et al., 2018) (resolution: 200 parcels per hemisphere) applied using mris_ca_train to the native white/gray boundary of the cortical surface (smoothwm) as generated using Freesurfer’s recon-all. Cerebellar regions of interest were identified using the Diedrichson probabilistic atlas of the human cerebellum (Diedrichsen et al., 2011, 2009), normalized to MNI152 atlas space. Vertex time courses were extracted using custom Matlab (Mathworks, Inc) scripts in conjunction with Neuroelf (https://neuroelf.net/). Time courses z-score normalized then averaged into a single time series per parcel.

### Connectivity estimation

Functional connectivity was estimated using coherence measures between different regions of interest (ROIs) in the cerebral and cerebellar cortices. Coherence measures were computed using the wavelet toolbox (Grinsted et al., 2004), completed in Matlab (Mathworks, Inc.). Coherence was calculated across discrete non-overlapping temporal windows (20 volumes, or 40 s) and averaged across frequencies corresponding to the task-relevant spectral content in fMRI BOLD connectivity, approximately 0.06–0.12 Hz (Sun et al., 2005, 2004). This calculation yielded 28 unique coherence matrices representing the strength of each pairwise connection across the atlas parcellation.

### Dynamic community detection

We employed dynamic community detection algorithms across 28 temporal layers of functional brain connectivity, to evaluate the modular organization of the brain before and after cerebellar stimulation. The algorithm optimizes a modularity function Q to distill complex connectivity matrices into a series of coarse clusters or communities of networks across time and was implemented using generalized Louvain algorithm and standard optimization procedures (for review see (Garcia et al., 2018; Mucha et al., 2010; Lima Dias Pinto et al., 2024)).

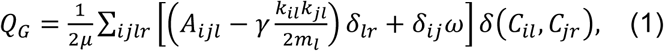

where, *Q_G_* is the generalized multilayer modularity function, indices *l* and *r* denote consecutive time layers (as shown in Figure 1), *A_ijl_* is the weighted edge between nodes *i* and *j* in layer *l*, *k_il_* is the degree of node *i* in layer *l*, *m*_*l*_ is the sum of the edge weights of layer *l*, *μ* is the sum of the edge weights of all time layers, *C_il_* is the community affiliation of node *i* in layer *l*, and *δ* is the Kronecker delta, which equals 1 if *i*=j, and 0 otherwise. The community assignments are dependent on two parameters: (1) a structural resolution γ parameter and (2) a temporal resolution ω parameter. We used γ=1 and ω =1 in this analysis. This, on an average, yielded 5 communities (SD = 0.9) that spanned the 18:40 minute scan for both the pre- and post- cerebellar stimulation conditions. Due to the heuristic nature of the generalized Louvain algorithm, we ran 100 iterations of community detection for every individual and condition (pre- and post-stimulation) and the estimated metrics (discussed below) were averaged across these iterations.

**Figure 1:**
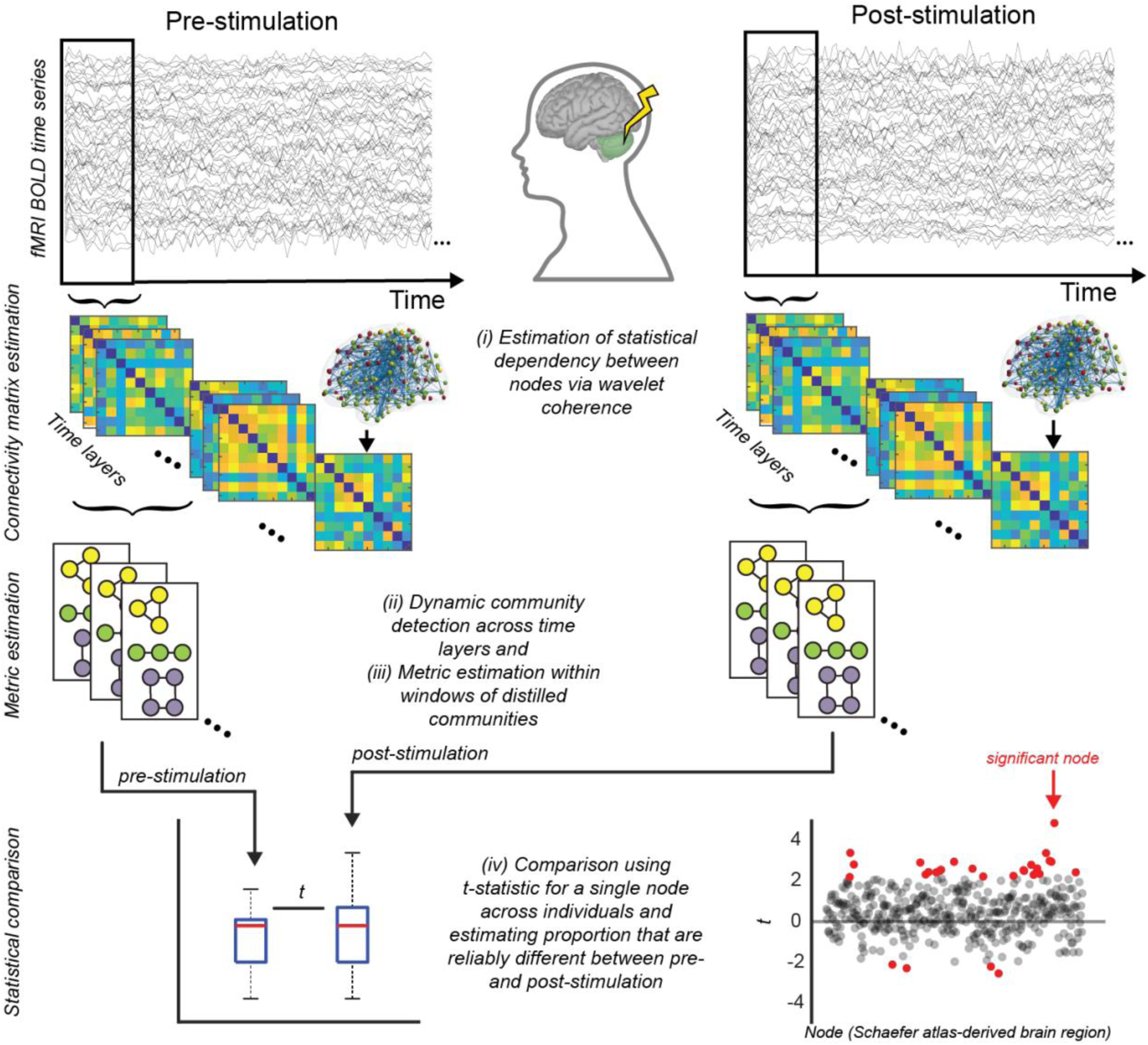
Schematic of the experimental and analysis design. Resting state fMRI activity was measured before and immediately following 20 minutes of 1 Hz inhibitory active stimulation (rTMS) delivered to the right cerebellum. Our analysis steps included: (i) the estimation of functional connectivity between 432 ROIs (400 cortical parcels spanning left and right hemispheres, and 32 cerebellar regions of interest) via wavelet coherence in non overlapping temporal windows (40 s each); (ii) dynamic community detection on the temporal connectivity matrices (time layers) to distill a series of community structures; (iii) extraction of complex network metrics quantifying network reconfigurations and (iv) statistical comparisons using the *pre-stimulation* condition as the baseline.

### Quantifying dynamic reconfigurations

In our study, we computed a range of network metrics to quantify the dynamic reconfigurations in cerebral cortex and cerebellar connectivity following cerebellar stimulation. Our metrics of interest fall into two broad categories. The first category includes metrics that are agnostic to any reference community structure: node allegiance, flexibility and promiscuity, estimating node-pair-wise and nodal dynamic changes in affiliation. The second category comprises metrics that are calculated relative to a reference community structure (see Consensus community below): integration, recruitment, participation coefficient, and within-module degree. Together, these metrics provide a comprehensive view of the dynamic community structure and the roles individual nodes play within this structure.

#### Node allegiance

We calculated allegiance matrices to assess the pairwise relationship of the brain regions. Allegiance accounts for the proportion of the total time a pair of nodes belongs to the same community, and is defined as:

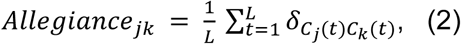

Here *L* is the total number of time layers, *C_l_*(*t*) denotes the community which contains the node *l* at time *t*, and *δ* denotes the Kronecker delta such that, 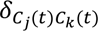 equals 1 if the nodes *j* and *k* are in the same community at time layer *t* and equals 0 otherwise.

#### Flexibility and promiscuity

Flexibility of a node is estimated by calculating the number of times a node changes its community allegiance normalized by the total number of allegiance changes possible (Bassett et al., n.d.). If *g_i_* is the total number of times node *i* changes its affiliation and *L* is the total number of time layers, the flexibility of node *i* is

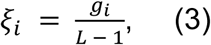

The promiscuity of a node is calculated as the fraction of all the communities in the network in which a node participated at least once (Papadopoulos et al., 2016). If a node participated in all the communities, its promiscuity score is 1.

#### Integration and recruitment

These metrics provide insight into the structure of observed network reconfigurations in relation to a reference network structure. The integration coefficient of a node (or region) corresponds to the average probability that this node shares allegiance - either transiently or sustained - with nodes from other reference-communities (Bassett et al., 2015; Mattar et al., 2015). The recruitment coefficient of a node corresponds to the average probability that this region is in the same network community as other regions from its own reference-community (Bassett et al., 2015; Mattar et al., 2015).

#### Participation coefficient and within module degree

The participation coefficient is a metric that quantifies the diversity of a node’s connections across different modules or reference-communities in a network (Guimerà and Nunes Amaral, 2005). For a node i, participation coefficient P_i_ is calculated as:

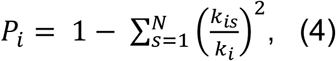

Where k_is_ is the number of connections node i has to the nodes in community s, k_i_ is the total degree of node i, and N is the number of communities. The participation coefficient ranges from 0 to 1 such that a value close to 0 indicates that the node’s connections are primarily within its own reference-community, while a value close to 1 suggests that the node has more uniformly distributed connections across multiple communities.

The within-module degree, often denoted as z, measures how well a node is connected within its own community or module. It is also calculated relative to a reference community structure and is defined as:

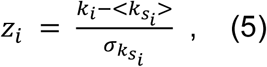

here, k_i_ is the number of links of node i to other nodes in its module 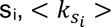 is the average of k over all the nodes in s_i_ and 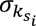 is the standard deviation of k in s_i_. A high z value indicates that the node is highly connected within its own community, serving as a potential hub or central node.

These metrics provide nuanced insights into the roles individual nodes play within their own communities and in the network at large. The participation coefficient helps in understanding a node’s role in intermodular connectivity, while the within-module degree focuses on intramodular connectivity. Here, we estimated these metrics using allegiance matrices to directly assess community-base reconfigurations.

#### Temporal variation in the metrics of interest

We computed the metrics of interest on resting state images collected pre- and post- cerebellar stimulation. Within a condition, using a sliding window approach with 90% overlap between adjacent estimates, we obtained community structure for 10 consecutive windows or temporal layers to estimate the metrics of interest, which led to 19 estimation points for the entire scan with each estimation point representing roughly 7 minutes of dynamics. This approach not only quantified the network reconfigurations, but also allowed us to probe where the maximum effect of stimulation lies and how it changes immediately after stimulation due to underlying neurological processes.

### Consensus community

Consensus communities in dynamic community detection refer to a technique that aims to identify stable or recurring patterns of community structure across multiple temporal windows or across different individuals. This is achieved by aggregating the community affiliations of nodes over time such that the modularity function across multiple layers (temporal or individual) is optimized to best represent the aggregated data. By focusing on consensus communities, the technique filters out transient or spurious community affiliations, thereby increasing the reliability of the detected community structure.

In this study we obtained a consensus community structure for each individual (Supplemental Fig. S2) by first calculating a consensus similarity across all the 28 temporal windows for the total duration of each scan. Consensus similarity estimates a single representative partition from a set of partitions (here 28) that is the most similar to all others. We then calculated the consensus iterative for all the iterations (i.e., 100 iterations) of dynamic community detection. Consensus iterative identifies a single representative partition from a set of partitions (here 100), based on statistical testing in comparison to a null model (Bassett et al., 2013).

Individually obtained consensus communities allow for the exploration of individual differences in community structure, which can be essential for understanding subject-specific responses or conditions. It allows for more nuanced inter-subject comparisons and can help in identifying subject-specific markers or predictors.

### Statistical analysis

We estimated the metrics of interest for pre- and post-stimulation conditions independently. We used paired t-tests with significance level 0.05 to draw statistical comparison of various metrics between conditions for the regions of interest (nodes).

## Results

In this study, we characterized how noninvasive inhibitory neuromodulation delivered to a region of the lateral cerebellum associated with cognitive function impacts network organization within the cerebral cortex. In brief, to accomplish this we measured intrinsic fMRI activity in the resting state from seventeen individuals before and immediately after they received 20 minutes of 1 Hz inhibitory active stimulation (rTMS) in the right cerebellum Crus I. Through a series of network dynamics analyses, we evaluated the overarching community structure, the dissimilarity in network nodal properties before and after stimulation, and the nature of the dynamic network changes following active stimulation, focusing on the roles of integration and recruitment. Additionally, we characterized the dynamic hub and integrator nodes throughout the cortex based on metrics like the participation coefficient and within-module degree. As described in Figure 1, our analysis scheme involved 5 steps: (1) atlas parcellation of BOLD activity, (2) functional connectivity estimation via wavelet coherence in non overlapping temporal windows, (3) dynamic community detection, distilling these functional connectivity matrices into a series of discrete communities, (4) dynamic metric calculation, and (5) statistical comparisons using the *pre-stimulation* condition as the baseline.

### Community structure and reference community

Given time evolving network connectivity, we first identified the community structure in networks during the entire scan using dynamic community detection (Garcia et al., 2018; Lima Dias Pinto et al., 2024; Mucha et al., 2010). We leveraged modularity maximization, which aims to identify network communities or modules such that within community connections are maximized and between community connections are minimized, and our analyses revealed a broad community structure that on an average consisted of 5 (SD = .9) communities within an individual for any given time window. This structure serves as the foundation for our subsequent analyses, providing a framework by which to understand the complex interplay between different regions of the brain.

### Dissimilarity before and after stimulation

Figure 2 quantifies the dynamic community changes across time, agnostic to a reference community. *Flexibility* broadly refers to the likelihood that a node changes its community affiliation across time and increased flexibility has been broadly linked to a variety of cognitive phenomena, including memory (Braun et al., 2015), learning (Bassett et al., n.d.), influence (Cooper et al., 2018), opinion change (Lima Dias Pinto et al., 2022), and has even been proposed to be a potential marker of cognitive flexibility (Xiao et al., 2022). As a complimentary metric to this generically dynamic metric, we also estimated *promiscuity* which quantifies the proportion of communities that a node will affiliate with at least once through the time period of question. Whereas *flexibility* characterizes the coarse changes of a node across time, *promiscuity* gives sharper insight into the type of affiliative changes a node may traverse.

**Figure 2:**
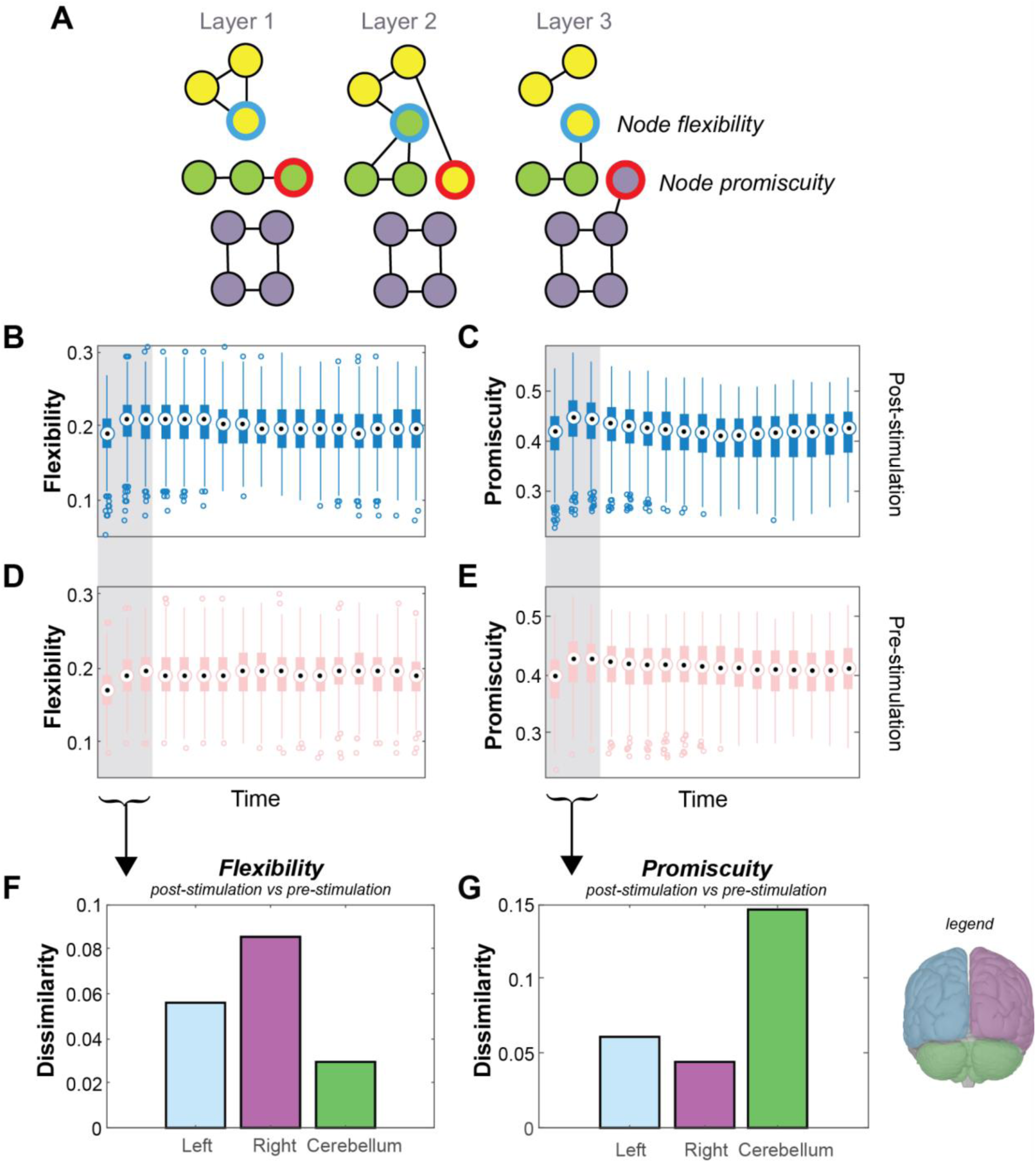
Impact of cerebellar stimulation on propensity of module or community reconfigurations. (A) We evaluated two metrics, flexibility and promiscuity, which assess the likelihood and heterogeneity of nodal allegiance changes within the estimated communities or modules. Temporal variation of flexibility and promiscuity (B)-(C) post cerebellar stimulation and (D)-(E) pre-stimulation resting state. (F)-(G) Statistical significance was computed using t-test comparisons between post-stimulation and pre-stimulation conditions using node-wise scoring, averaged over the first three windows to capture the immediate effects of the stimulation that are known to decay over time. Dissimilarity represents the proportion of nodes within each region (left and right cerebral cortex, and cerebellar cortex) for which flexibility or promiscuity differed significantly between post- and pre-stimulation conditions, using t-tests (p-value < 0.05, uncorrected).

To quantify the effect of cerebellar stimulation on network organization, we calculated these metrics in discrete windows across time for both pre- and post-stimulation conditions (Figure 2B-E). The “dissimilarity” score reflects the proportion of paired t-tests on nodal properties that are significantly different before and after stimulation. Considering the known decay of the effect of rTMS across time (Chen et al., 1997; Edwards et al., 2019; Pascual-Leone et al., 1998) and previous work that has shown network changes to dissipate within 15-20 minutes of rTMS in a similar protocol (Garcia et al., 2020b; Muellbacher et al., 2000), we focused the dissimilarity analysis on the metrics derived from the first 8 minutes (240 volumes) to understand the cortical effects of cerebellar stimulation (also see Supplemental Figure S1). The first 3 time windows show an overall dissimilarity of 0.07 in flexibility and 0.06 in promiscuity, scores that are better than by chance with a traditional statistical significance level of *p* = .05. A closer inspection of the dissimilarity distributions within the brain reveals more striking differences in dissimilarity when comparing the dynamic metrics flexibility and promiscuity in the left and right cerebral hemispheres and the cerebellum (Figure 2F-G). Notably, for flexibility, the right cerebral cortical regions has three times the proportion of flexibility changes to nodes as compared to the cerebellum (0.09 vs 0.03), indicating that after cerebellum stimulation, many right cortical regions change their affiliation. For promiscuity, however, the dissimilarity score for the cerebellum (∼ .15) is nearly three times that of the left and right (∼ .05) cortical regions, indicating that when the cerebellar nodes change their community affiliation after stimulation they are more likely to do so indiscriminately.

We next inspected the nature of these changes indicated by the dissimilarity metric. Figure 3A-B displays the direction of these flexibility and promiscuity changes, with the preponderance of nodes demonstrating an increase in these metrics following cerebellar stimulation. Among the nodes impacted by stimulation, 68% and 69% increased in flexibility and promiscuity, respectively, with only 2-4 cortical nodes showing a significant decrease in either metric. Inspecting the spatial distribution of these stimulation induced network dynamics revealed the nodes that significantly differed in their flexibility and promiscuity were not localized to a particular part of the brain; instead, they were distributed throughout the cerebral cortex, evidence for a more global response to cerebellar stimulation, at least cortically.

**Figure 3:**
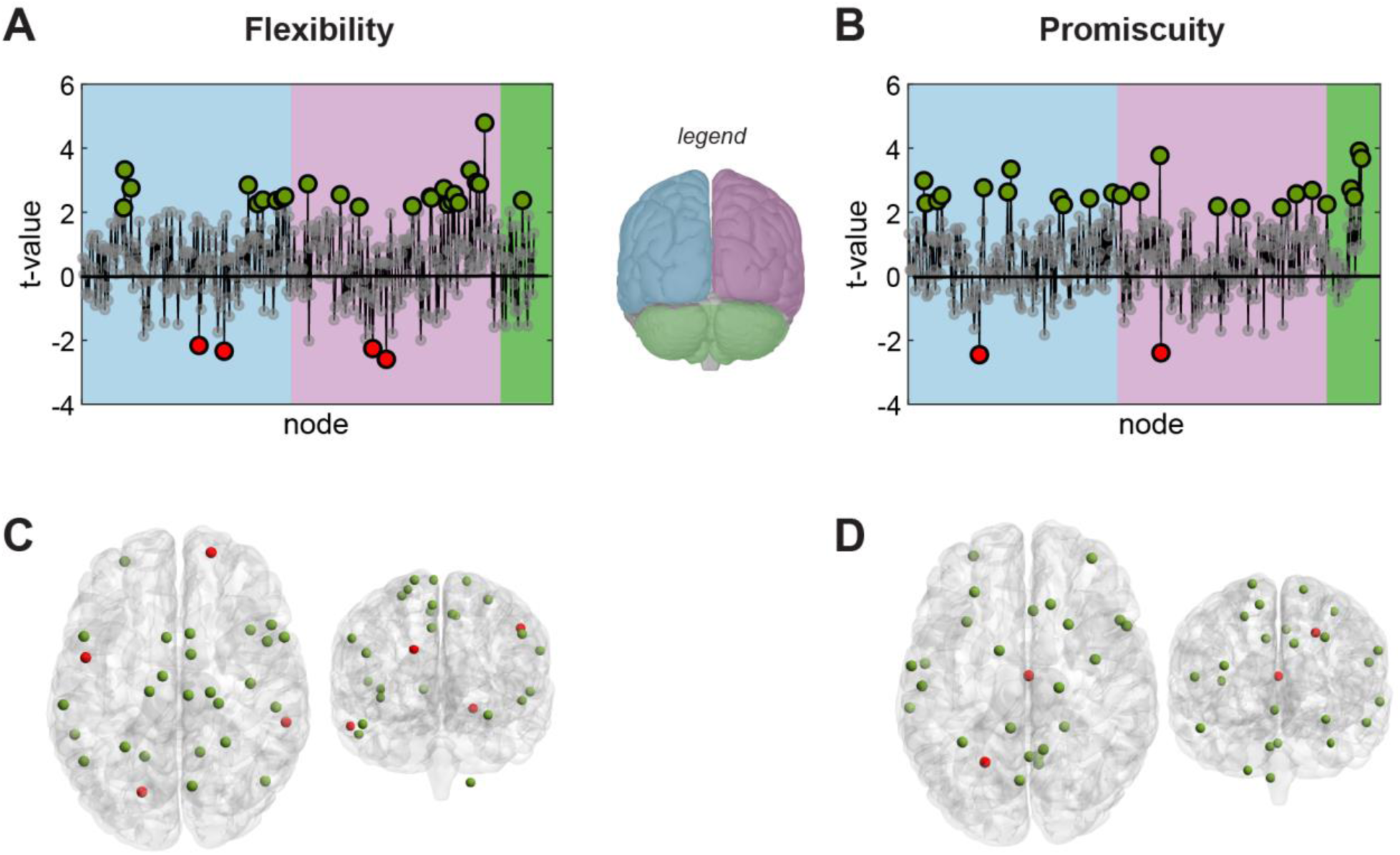
Flexibility and promiscuity increase post cerebellar stimulation. (A)-(B) Estimated t-values when flexibility and promiscuity were compared between post-stimulation and pre-stimulation conditions. Positive t-values represent higher flexibility or promiscuity post stimulation. Larger dots represent those nodes with significant differences (p-value < 0.05, uncorrected). (C)-(D) Distribution of nodes with significantly different flexibility or promiscuity post stimulation. Green nodes significantly increased metric dynamics post stimulation.

### Dissimilarity in integration and recruitment due to cerebellar stimulation

While promiscuity and flexibility can capture the changes of community affiliation across time, they do so in a very generic manner with no information on the specificity of these changes within the community structure. For this reason, we next inspect two other, reference-community specific metrics that may capture potentially nuanced changes. Node *integration* refers to the average probability that a node is in the same community as the nodes from other reference communities, and *recruitment* refers to the average probability that a node is in the same community as other nodes from its own reference community (Figure 4A).

**Figure 4:**
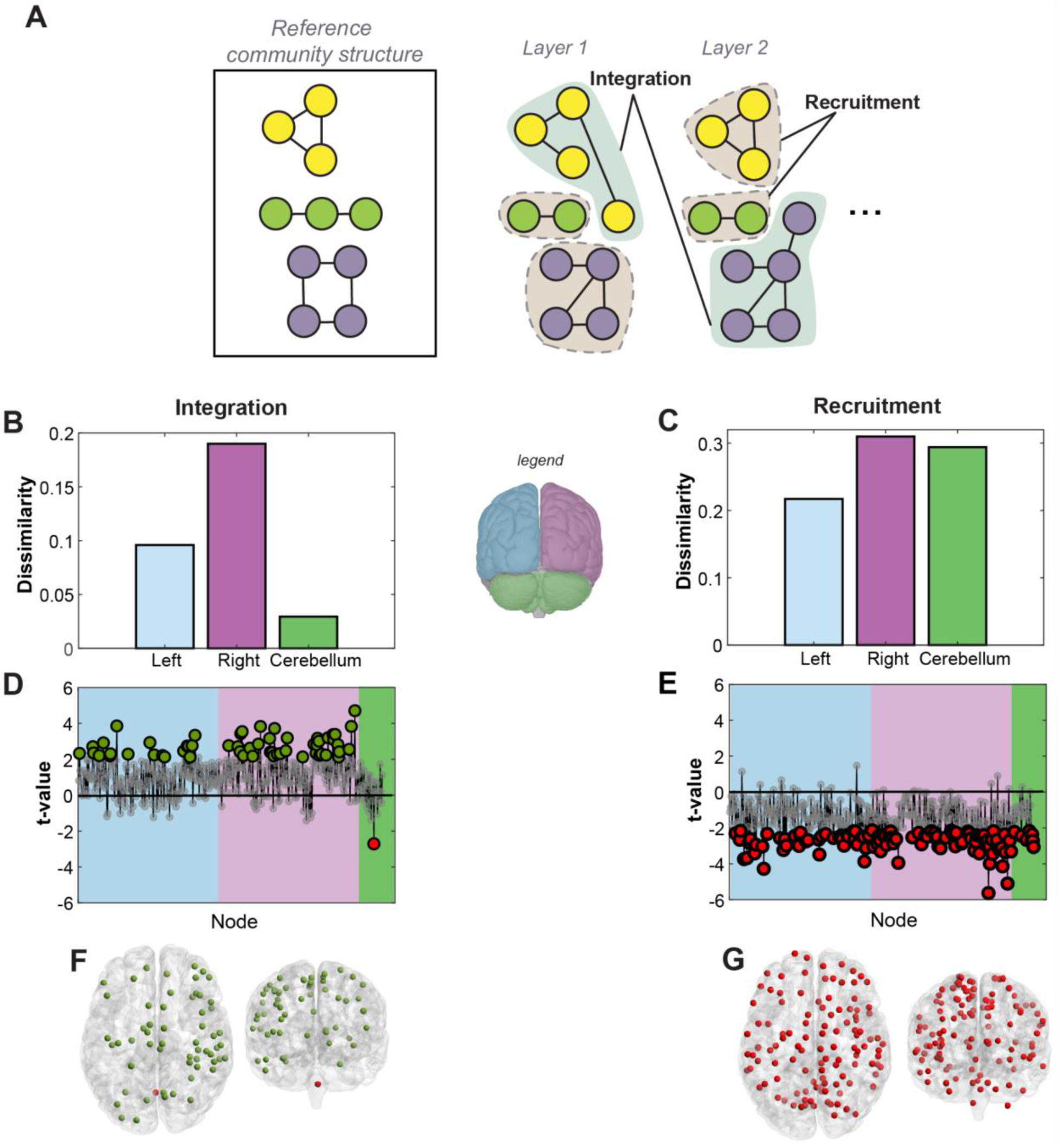
Community structure changes due to cerebellar stimulation. (A) Integration and recruitment quantify changes in node allegiances with respect to a reference community structure. Integration quantifies the probability of a node to be in the same community as the nodes from other reference communities and recruitment quantifies the probability of a node to be in the same community as the nodes from its own reference community. (B)-(C) Dissimilarity represents the ratio of nodes within each region (left, right, and cerebellum) for which integration or recruitment was significantly different between post-stimulation and pre-stimulation conditions, using t-tests (p-value < 0.05, uncorrected). (D)-(E) Estimated t-values. Positive t-values represent higher integration or recruitment post-stimulation. Larger dots represent significant differences. (F)-(G) Distribution of nodes with significantly different integration or recruitment post stimulation. Green implies an increase. Large proportions of nodes display significant changes in integration and recruitment which we attribute to a stimulation driven change in overall community structure.

We used the community structure in the pre-stimulation condition as a baseline modal community structure against which to compare changes due to cerebellar stimulation. We estimated the modal community structure across both time and subjects using a *consensus community* estimation, which constructs the most representative partition across time and iteration using consensus similarity and consensus iterative processes (Bassett et al., 2013). On average across subjects, 10 (SD = 4.5) communities were found when assessing the community structure within an individual subject across time and iterations. As visualized in the consensus community structure in the resting state data acquired prior to stimulation (Supplemental Figure S2), we observed highly variable architectures when assessed across subjects. Thus, for these network metrics that are calculated relative to a *reference* community, we used the within-subject consensus community structure, as our data indicates this may best capture the dynamic changes of not only cerebellar stimulation but also best characterize the *resting state* functional architectures within each individual (Supplemental Figure S3).

We find that many cortical regions exhibited high levels of dissimilarity from the consensus community structure following stimulation. Approximately 20% of nodes in the right cortical hemisphere and approximately half of that (10%) in the left cortical hemisphere showed broad differences in integration following stimulation, with little change observed in the cerebellum nodes. Conversely, a high proportion of nodes in both the right hemisphere and the cerebellum (∼ 30%) showed changes in recruitment following stimulation, with slightly fewer in the left hemisphere.

As before, we next inspected the direction of this dissimilarity. Interestingly, cortical nodes with significant changes in integration scores induced by rTMS illustrated an increase in integrative dynamics, evidence that neuromodulatory effects in cortical dynamics were distributed across networks. Likewise, all nodes with significant changes in recruitment scores following stimulation had lower scores of network recruitment, further evidence that neuromodulation over the cerebellum induced a fractionation in the existing community structure.

### Dynamic characteristics of the cortical and cerebellar nodes

Finally, to understand the impact of cerebellar stimulation, we assessed the network properties of the cerebellar nodes by analyzing the *participation coefficient* and *within-module degree* in baseline (pre-stimulation) condition and comparing them to nodes in the cerebral cortex. In brief, the participation coefficient estimates the distribution of the edges (allegiances), where a high value would suggest distributed connections within different communities. Within-module degree on the other hand quantifies how connected a node may be to its own reference community. Thus, these two metrics may then inform whether nodes may be “dynamic hubs” (high within module degree) or “dynamic integrators” (high participation coefficient and low within module degree).

Note that this analysis is analogous to previous examinations of hubs and integrators using these metrics (e.g., (Guimerà and Nunes Amaral, 2005)), except for two things: first, rather than focusing on direct functional connectivity estimates (i.e., wavelet coherence), our analyses were defined on *allegiance*, the pairwise estimate that a node-pair appears in the same community across time. We therefore estimated these metrics on dynamic network measurements rather than static. Dynamic network measurements also capture the evolving connectivity patterns, revealing regions that act as integrators by flexibly connecting with various communities or modules across different time points that cannot be revealed using static network measurements. Second, instead of using a predefined functional system as reference communities, we utilized individualized *consensus community* structures to calculate the network metrics allowing the *functional* networks to change based on the natural neural communication patterns within an individual. This individualized dynamic perspective highlights the node’s role in coordinating and integrating information across the brain, adapting to the shifting demands of cognitive and motor tasks for each individual.

Interestingly, this analysis revealed that all the dynamic hubs were located in the cortical regions, while the dynamic integrators were overwhelmingly represented by cerebellum nodes. Inspection of the summary bar plots in Figure 5, we see that nearly 2 times the proportion of cerebellum nodes, compared to cortical nodes, are likely to be dynamic integrators, as defined by being in the top 50 percentile of participation coefficients and below median within-module degrees. Using the 95th percentile as a cut-off for being a dynamic hub, no cerebellum node reached this threshold. To further clarify this distinction, we also examined the allegiance changes across the cerebellum and cortex, investigating whether the dynamic reconfigurations induced by the stimulation may be driven by a prioritization of dynamic changes through cortical hubs. Instead, we see that the allegiance changes in the cerebellum are on par with those in the cerebral cortex, suggesting that the dynamic changes are likely being captured solely by the integration and recruitment changes (see Supplemental Figure 4).

**Figure 5:**
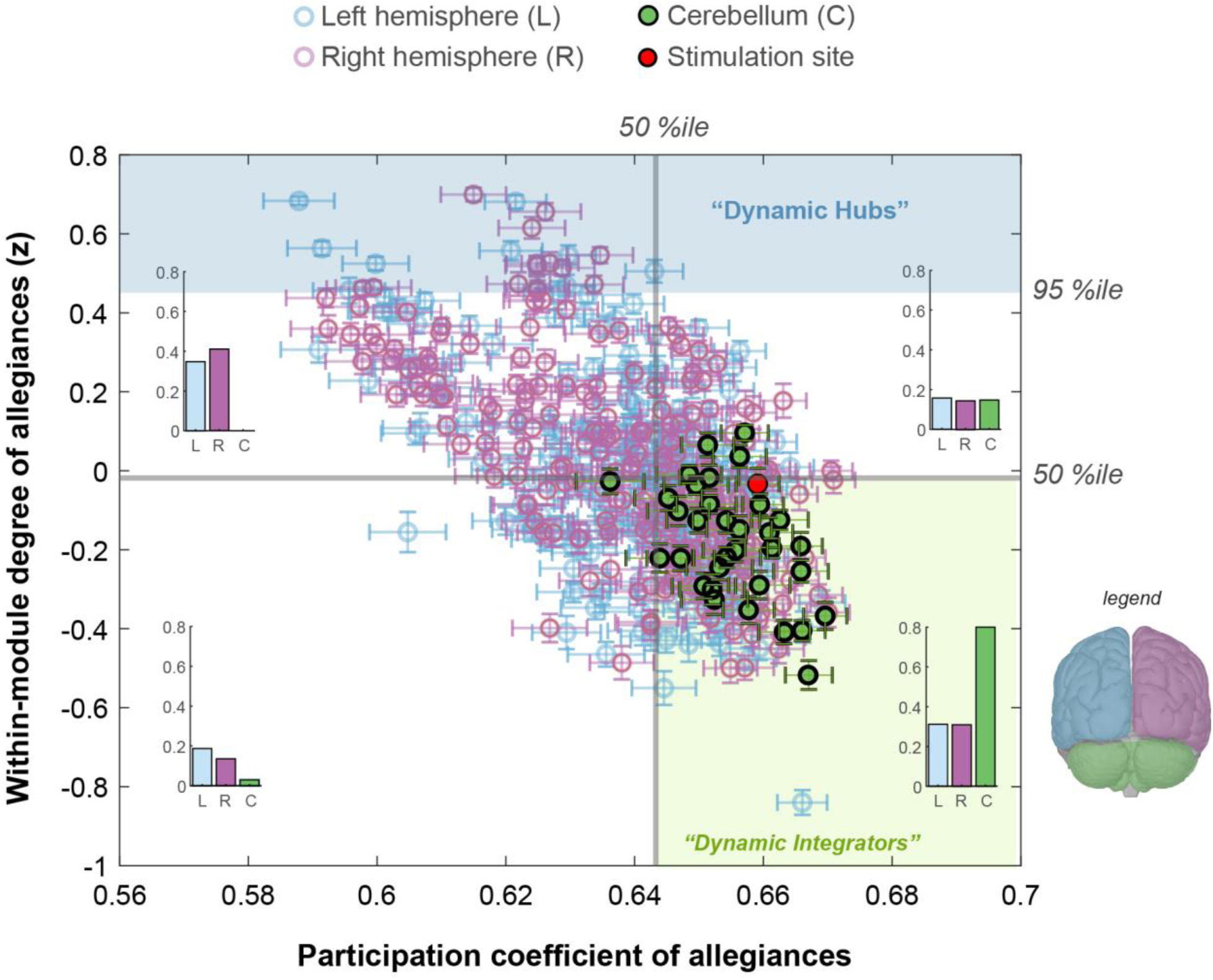
Dynamic characteristics of cortical and cerebellar nodes. Participation coefficient and within-module degree estimated using allegiance matrices for baseline (pre-stimulation) condition. Here, the participation coefficient denotes the likelihood for a node to have allegiances with nodes outside of its reference community; and the within-module degree represents the tendency of a node to have allegiances with the nodes from its own reference community. Nodes with very high within module degree (above 95%ile) are identified as dynamic hubs. None of the cerebellar nodes fall into this category. Nodes with high participation coefficient (above 50%ile) and low within module degree (below 50%ile) are identified as dynamic integrators, which primarily make allegiances with nodes outside of their reference community. Majority of cerebellar nodes fall into this category. For each quadrant (divided by 50%ile lines) we show the proportion of cortical (left and right) and cerebellar nodes using inset bar plots.

## Discussion

Neuromodulation has the potential to induce prolonged intervals of altered cortical dynamics between functionally connected circuits (Edwards et al., 2019; To et al., 2018). In this study we have characterized the dynamics of resting state community structure following 1 Hz inhibitory stimulation (“perturb and measure”) over Crus 1 of the lateral cerebellum. The neuromodulation induced significant changes in node flexibility and promiscuity among nodal affiliations in both cerebral and cerebellar cortex, indicating that the cerebellar stimulation enhanced intrinsic network flexibility and resulted in a period of rapid reconfigurations. Interestingly, we found that the specifics of these changes could be captured by the integrative and recruiting behaviors of cerebellar nodes. Consistent with the role of the cerebellum as a mechanism for promoting efficiency in cognitive and motor function via the coordination of neural units (Ito, 2008; Koziol et al., 2014; Schmahmann, 2010), cerebellar nodes emerged as dynamic integrators within the brain, in contrast to the dynamic hubs found in cortical regions.

Our findings illustrate the integrative nature of the cerebellum and the potential for widespread disruption of functional architectures when that integrative function is disturbed, similar to maladaptive spreading of dysfunction via integrator nodes in the case of disease (Fornito et al., 2015). Our approach further highlights the nuanced role of network dynamics in understanding the interplay of large-scale functional systems.

### Cerebellar stimulation increases network flexibility

Intrinsic network structure is not static, but instead changes dynamically over time during cognitive tasks (Telesford et al., 2016) and at rest (Dong et al., 2019; Haller et al., 2013; Rungratsameetaweemana et al., 2022; Wang et al., 2020). We evaluated two key metrics of dynamic network behavior that characterize patterns in community affiliation over time, with emphasis on the movements and distribution of nodes across community boundaries.

Flexibility characterizes the likelihood of nodal affiliation changes over time, with studies showing that this metric correlates with improved learning rates (Bassett et al., 2011), increased reinforcement learning of visual cues and outcomes (Gerraty et al., 2018), improved working memory performance (Braun et al., 2015), the need for cognition and creative achievement (He et al., 2019), and the resistance to online influence (Lima Dias Pinto et al., 2022). We found that a significant minority of nodes, particularly in the right cerebral cortex, have increased flexibility in the brief interval following inhibitory stimulation, consistent with previous demonstrations that neuromodulation over the cerebellum propagates to functionally connected circuits in the cerebral cortex (Farzan et al., 2016; Halko et al., 2014; Rastogi et al., 2017). That cerebellar rTMS increased cortical flexibility is further evidence for the potential of neuromodulation to induce functional plasticity into network architectures that, rather than being purely disruptive, may allow for meaningful and structured dynamic reconfigurations of community structure.

Promiscuity characterizes the distribution of these reconfigurations across potential unique affiliations, computed as the likelihood of nodes affiliating with every other community at least once. We found that, on average, 40% of the nodes *increased* promiscuity after stimulation (Figure 3B), with cerebellar nodes more likely to exhibit distributed dynamic changes that include affiliations to many other communities. This finding is consistent with the characterization of inhibitory rTMS as inducing a disruption in underlying cortical dynamics (Edwards et al., 2019; Siebner et al., 2022), and further demonstrates substantial and global cortical response to cerebellar stimulation. Importantly, nearly all of the nodes that changed in flexibility, exhibited an *increase* in flexibility (68%).

At first blush and considering our findings, one may propose cerebellar stimulation as an intervention technique that may be harnessed and deployed for improved behavioral outcomes (e.g., enhanced working memory); however, as Safron et al. (Safron et al., 2022) point out, while elevated flexibility may be associated with adaptive behaviors, the literature appears to suggest that excessive flexibility could also reflect deleterious cognitive outcomes (Betzel et al., 2017) and could also be indicative of pathology (e.g., (Braun et al., 2016)). Moreover, within development, excessive flexibility in the visual areas of infants was negatively correlated with the rate of developmental milestone achievement (Yin et al., 2020), whereas increased flexibility in somatomotor areas and higher-order brain regions was typical during developmental progression, indicating a role for flexibility in adapting to novel situations and challenges.

It is important to note that the stimulation protocol employed in this study was low frequency repetitive stimulation, an inhibitory protocol. Behavioral outcomes using similar inhibitory protocols over the cerebellum have translated into poorer task performance, illustrated by an increase in error rates in a behavioral inhibition task (Esterman et al., 2017) and a decrease in sensory responses to auditory stimuli (Andrew, 2020). Additionally, previous studies utilizing cortical stimulation have documented relatively modest changes in flexibility following visual and parietal stimulation (Garcia et al., 2020a) with equally modest behavioral changes (Garcia et al., 2020b). Together, these results indicate the need for further research to evaluate the interaction between stimulation protocols and sites, and the potential to induce optimal levels of cortical network flexibility for particular tasks. The relatively large and non-specific changes in flexibility (and promiscuity) observed here in the resting state should be taken as evidence for the potential of cerebellar stimulation to elicit *excessive flexibility* in the cortex, with future research evaluating the specificity of behavioral outcomes.

### Cerebellar nodes uniquely act as integrators within the brain

In the human, Crus I and II of the lateral cerebellum are the largest of the lobes and are expanded in humans and apes relative to other mammals (Buckner, 2013; Habas, 2021; Strick et al., 2009). Resting state connectivity reveals these lobes to be nestled between and distinct from somatomotor maps in the cerebellum, and highly connected to prefrontal and parietal cortex (Buckner et al., 2011; Krienen and Buckner, 2009). Moreover, graphical analysis of the cerebellum in the resting state reveals its functional connections to have small world properties, meaning that the architecture of the graphs strike a balance between regular connectedness and fully random (Pezoulas et al., 2017), with significant individual differences in the small worldness and hierarchy (Chen et al., 2022). Consistent with that characterization, our supplementary analysis (Supplemental Figure 4) reveals that the allegiance patterns between nodes within cortex and cerebellum are quite non-specific. In other words, we observe that cerebellar integrative properties are not driven by clear hub communication to the cortex; instead, its connections are sparse and distributed across the brain.

Our final analysis focused on the flexible changes within networks and inspected the integrating and recruiting features of the changes within the cerebellum to the rest of the brain. Specifically, our analysis revealed approximately 30% of cortical nodes showed significant increases in integration and 50% of cortical nodes showed significant decreases in recruitment, and 30% of cerebellum nodes displayed a significant decrease in recruitment, compared to a single node that showed a decrease in integration. To further explore this finding, we next compared the within-module degree and participation coefficient within allegiance matrices estimating the pair-wise community changes within a reference community, essentially allowing nodes to be identified as *dynamic hubs* and *dynamic integrators* within the larger population. Interestingly, nearly all of the cerebellum nodes (80%) were dynamic integrators and only cortical regions were dynamic hubs. Overall, the findings reveal a nuanced picture of cerebellar functioning, characterized by distinct roles in integration and recruitment compared to cortical regions. Together, these unique integrative and recruitment properties may be intrinsically linked to the cytoarchitecture of the cerebellum, as a closed-loop highly dense computational module, where these emergent properties may be a direct result of this distinct cytoarchitectonic feature.

The concept of dynamic hubs and integrators further accentuates the cerebellum’s unique position within the brain’s functional network. The cerebellar nodes, characterized by their high participation coefficients and lower within-module degree, suggest a propensity towards integrating information across different brain regions. This is particularly intriguing given the emerging understanding that the cerebellum functions, at least in part, to optimize complex cognitive and motor functions (Koziol et al., 2014). Whereas damage to the cerebellum rarely results in profound cognitive deficits, neuropsychological tests reveal a more subtle but pervasive impact on language, working memory and executive function (Schmahmann, 2010). In both cognitive and motor function, the cerebellum is believed to support the integration of internal models (motor execution or mentalizing processes) within incoming sensorimotor signals so as to rapidly correct for errors and promote the skillful execution of cognitive and motor acts (Buckner, 2013; Ito, 2008). It should not be surprising, then, that our analyses of intrinsic connectivity reveals highly integrative properties in cerebellar nodes. Because these integrative properties are highly susceptible to neurostimulation, this indicates that the network roles reflect functional adaptability over structural constraints, suggesting a higher complexity in cognition and adding to our mechanistic understanding of how the cerebellum impacts broader cognitive and integrative functions.

### Cerebellar nodes’ non-specific integration

Further inspecting the connections of the highly integrative nodes of the cerebellum to the cortex, it may be efficient to utilize the hubs within cortex to maximally impact cortical functions; however, as shown in a supplementary analysis (Supplemental Figure S4), our results indicate that the allegiance patterns between nodes within cortex and cerebellum are quite non-specific. In other words, we observe that the cerebellum integrative properties are not driven by clear hub communication to the cortex; instead, its connections are sparse and distributed across the brain. This is similar to the concept of a *diverse club*, where nodes and edges are critical for efficient global communication (Bertolero et al., 2017).

Our dynamic augmentation to the diverse club concept, allows us to expand upon this construct, suggesting that dynamic diverse clubs within the brain may drive global communication and impact cognition in a general, in nonspecific fashion. In other words, by not focusing solely on cortical hubs, the cerebellum might gain a more comprehensive understanding of the overall *state* of the cortex, rather than simple read-out of dynamic hubs within cortex. Abstracting and integrating information from other cortical regions, the cerebellum may be integrating *contextual* information from many diffuse and functionally diverse cortical areas. This broad distribution of dynamic reconfigurations could allow the cerebellum to assimilate the rich orchestration of neural activity, enhancing its capacity to modulate and fine-tune cognitive and motor functions, across many spatial scales. Such an integrative role might enable the cerebellum to better predict and respond to the dynamic demands of the brain, maintaining a seamless coordination between different functional domains. This capacity for extensive contextual integration suggests that the cerebellum is not merely a passive recipient of cortical inputs but an active participant in shaping the brain’s functional architecture.

### Rapid and widespread dynamic whole-brain reconfigurations

In addition to the cerebellum specific results, our analyses using dynamic community detection revealed a time-evolving network connectivity with an average of five communities identified within an individual for any given time window. When comparing these networks to the pre-stimulation condition baseline, we observed highly variable community structures across participants, indicating a broad and dynamic architecture. Specifically, the consensus community estimation across subjects and time indicated poor alignment with standardized functional networks, such as the Default Mode Network and Control Network, with normalized mutual information (NMI) values never exceeding a moderate level of consistency (MI < 0.32). This variability and lack of alignment highlight the brain’s inherent elasticity and dynamic propensity (Safron et al., 2022). The observed poor alignment across individuals suggests that each individual’s brain network configuration is unique and constantly reconfiguring. This high level of plasticity underscores the importance of treating the brain as a highly dynamic system, capable of substantial reorganization in response to different stimuli or conditions. It implies that personalized approaches may be necessary for understanding and effectively targeting brain function and disorders, rather than relying on a one-size-fits-all model based on group averages (Dubois and Adolphs, 2016). The brain’s capacity for such extensive reconfiguration may be fundamental to its adaptability and resilience, allowing for a flexible response to ever-changing environmental demands.

## Conclusion

Our findings suggest that the cerebellum, through its integrative properties and sparse, distributed connections to the cortex, plays a unique role in modulating brain network dynamics. Unlike the cortical regions that act as dynamic hubs, the cerebellum emerges as a dynamic integrator, facilitating the rapid and flexible reconfiguration of brain networks. This broad connectivity pattern may provide the cerebellum with a more comprehensive understanding of the cortical state, integrating rich contextual information that enhances its ability to modulate cognitive and motor functions. This integrative capacity could provide a mechanism for the dysmetria of thought, where cerebellar disruption can impair higher-order behavior by affecting the cerebellum’s modulation of cognitive operations. Extending this more generally, we posit that the cerebellum could play a role in navigating cognitive maps (e.g., (Whittington et al., 2022)) rapidly and effortlessly, as proposed by recent theories, underscoring its potential as a critical element in intelligence (Hawkins et al., 2019). This perspective could suggest avenues for future research, exploring the cerebellum’s contributions to brain function, perhaps encompassing a fundamental role in cognitive integration and adaptability (Honda et al., 2018). The brain’s inherent plasticity and dynamic nature, highlighted by our study, further reinforce the need to treat it as a flexible system, capable of extensive reorganization in response to varying demands, and underscores the potential for personalized approaches in neuroscience research and therapeutic interventions.

## Acknowledgements

This research was supported by the U.S. Army DEVCOM Army Research Laboratory through mission funding (JOG), army educational outreach program (KB, W911SR-15-2-0001) and the Italian Ministry of University and Research (ZC; PRIN 20203LT7H3). The views and conclusions contained in this document are those of the authors and should not be interpreted as representing the official policies, either expressed or implied, of the US DEVCOM Army Research Laboratory or the U.S. Government. The U.S. Government is authorized to reproduce and distribute reprints for Government purposes notwithstanding any copyright notation herein.

## Author contributions

KB: Conceptualization, Formal analysis, Methodology, Software, Visualization, Writing – original draft, Writing – review & editing

ZC: Funding Acquisition, Investigation, Writing – review & editing

VO: Investigation, Writing – review & editing

CF: Investigation, Writing – review & editing

EDG: Data curation, Writing – original draft, Writing – review & editing

JOG: Conceptualization, Methodology, Software, Visualization, Funding Acquisition, Writing – original draft, Writing – review & editing

## Supplementary Material

**Figure S1:**
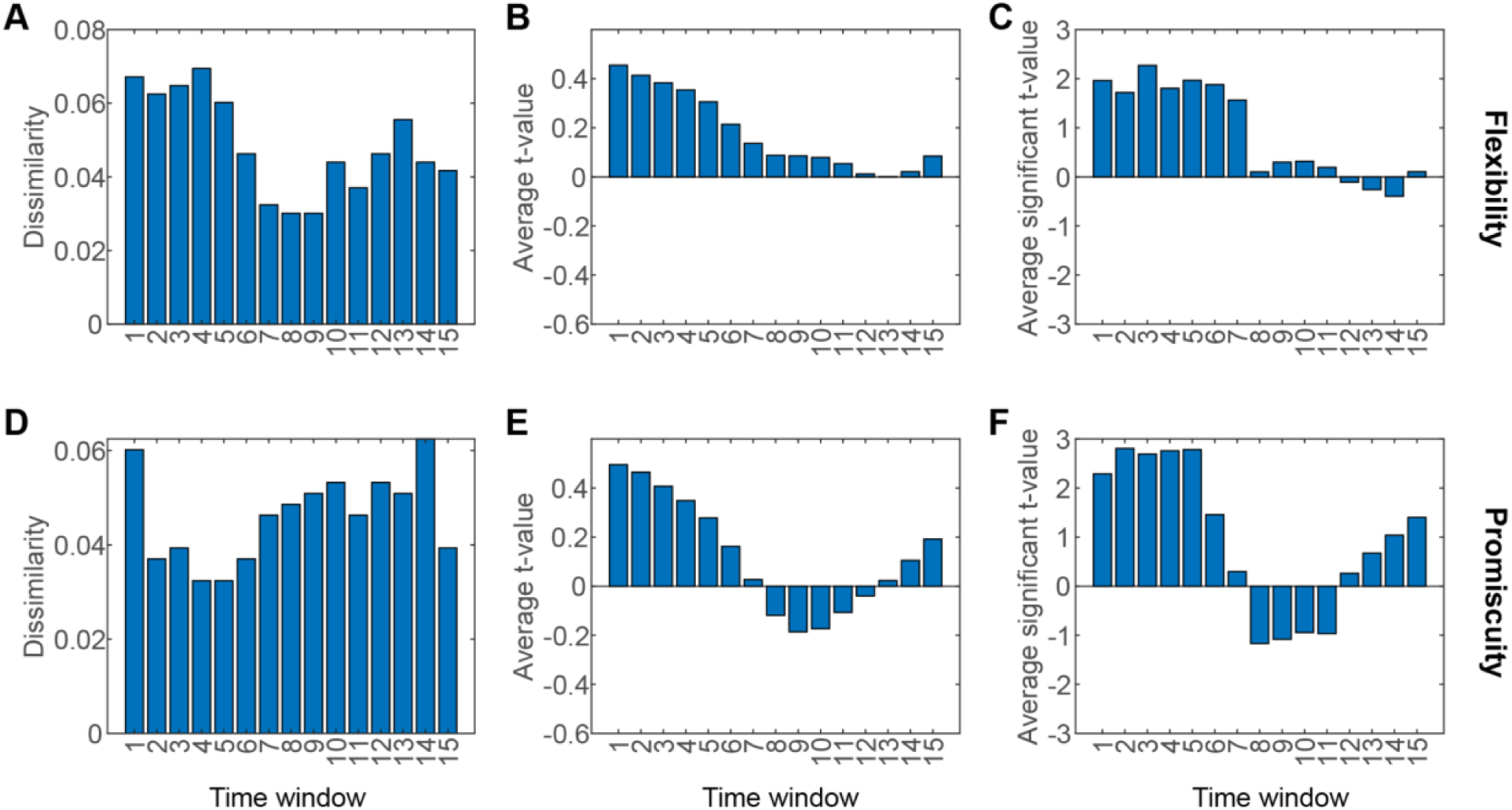
Comparing post-stimulation and pre-stimulation conditions as a function of time. We used sliding time windows with 67% overlap. Each time window here represents an average of three consecutive time points for which metrics of interest were calculated (e.g., in Figure 2B-E). (A),(D) Dissimilarity as a function of time for flexibility and promiscuity respectively. (B),(E) Average t-value as a function of time across brain nodes when flexibility and promiscuity were compared post-stimulation as compared to pre-stimulation. High positive values indicate strong increase in the metrics post-stimulation across many nodes. Values close to zero indicate either low t-values across all nodes or symmetrically distributed positive and negative t-values, likely indicating the fading impact of stimulation. (C),(F) Average t-value only for nodes that showed significantly different (p-value < 0.05, uncorrected) flexibility and promiscuity respectively, post-stimulation compared to pre-stimulation.

**Figure S2:**
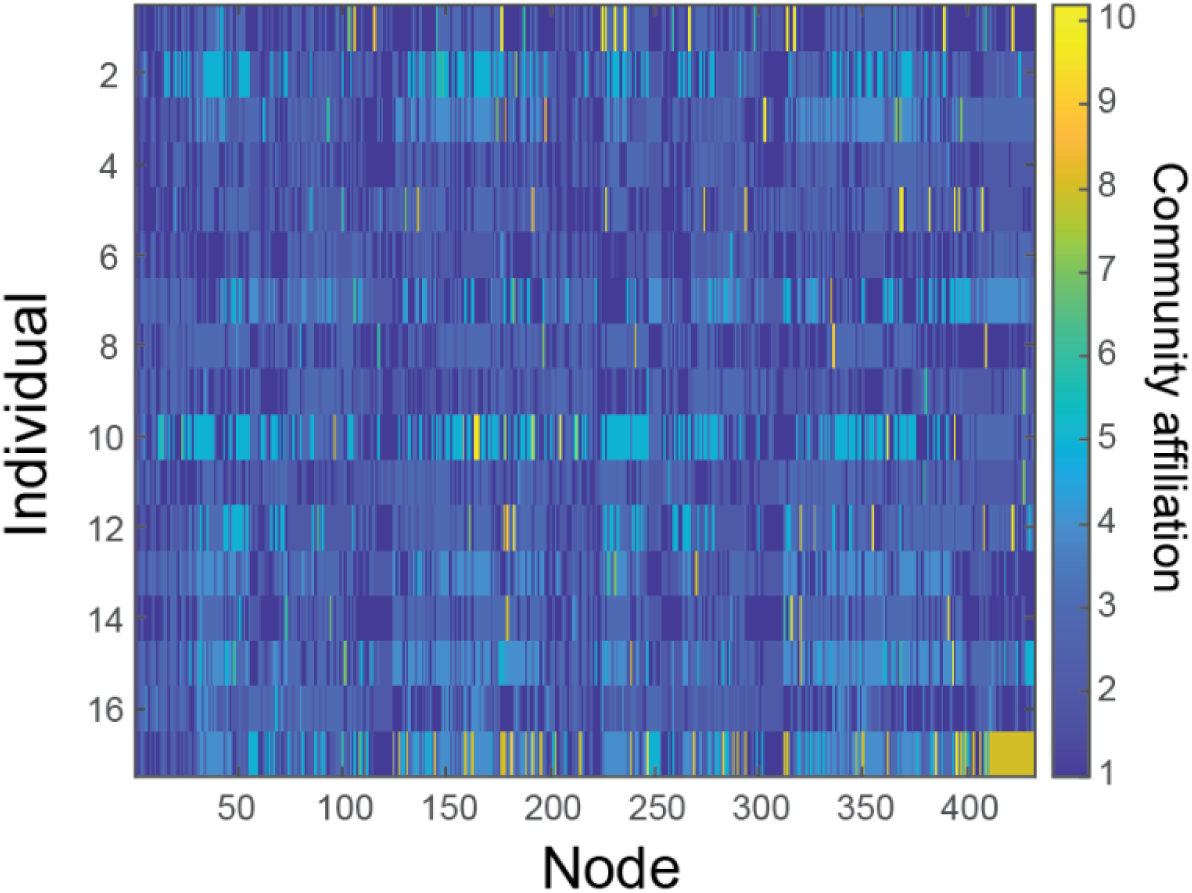
Individual consensus community. We obtained a consensus community structure for each individual by first calculating a consensus similarity across all the temporal windows for which community detection was performed (i.e., 28 windows) and then calculating consensus iterative for all the iterations (i.e., 100 iterations) of dynamic community detection. Consensus similarity estimates a single representative partition from a set of partitions (here 28) that is the most similar to all others. Consensus iterative identifies a single representative partition from a set of partitions (here 100), based on statistical testing in comparison to a null model. Occasionally, in consensus community structure we observed communities with single nodes, which were excluded from further analysis. Most of the nodes were distributed between 6 or fewer communities in any given subject.

**Figure S3:**
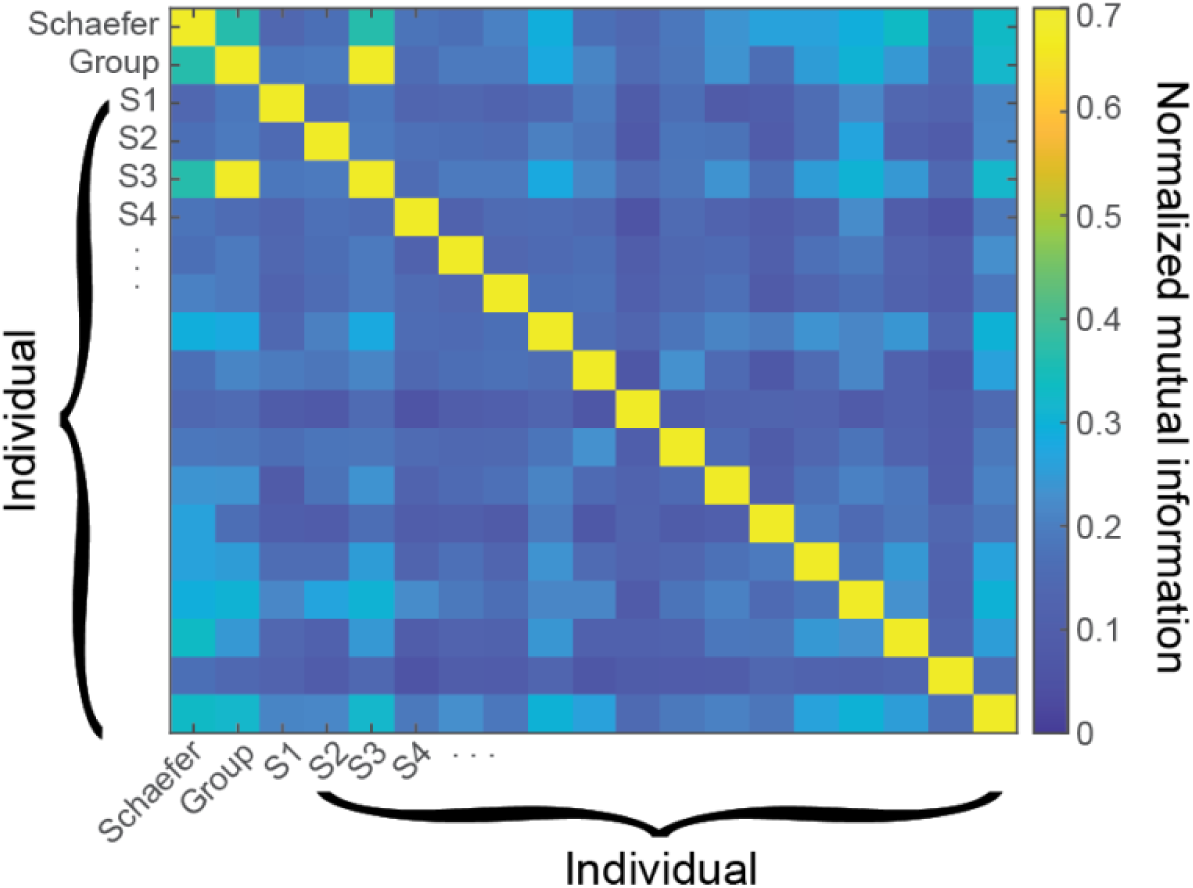
Comparison between different community partitions using normalized mutual information (NMI) between different community structures. NMI is a measure of similarity between two different partitions of community structure and is bounded between 0 and 1. We used two different functional characterizations of *standardized networks.* The first is a data driven community structure that is completed by estimating the consensus similarity between individual consensus communities (from Figure S1), which found 10 communities across subjects (“Group”), and another using the functional atlas labels provided by the chosen parcellation, which used labels consistent with previously understood functional networks (e.g., Default Mode Network, Control Network, Limbic System, etc.) as part of the Schaefer parcellation. We estimated NMI between communities, comparing each subject to one another in addition to the *Group* network and *Schaefer* network compositions. Neither consensus community *Group* network nor *Schaefer* network compositions substantially deviated from the inter-subject NMI, where the average NMI on inter-subject NMI was M = 0.16 (SD = 0.03), and that of the Group was 0.21 and Schaeffer was 0.23. In fact, NMI across all comparisons never exceeded 0.32, which indicates a poor alignment of overall network architecture across individuals.

**Figure S4:**
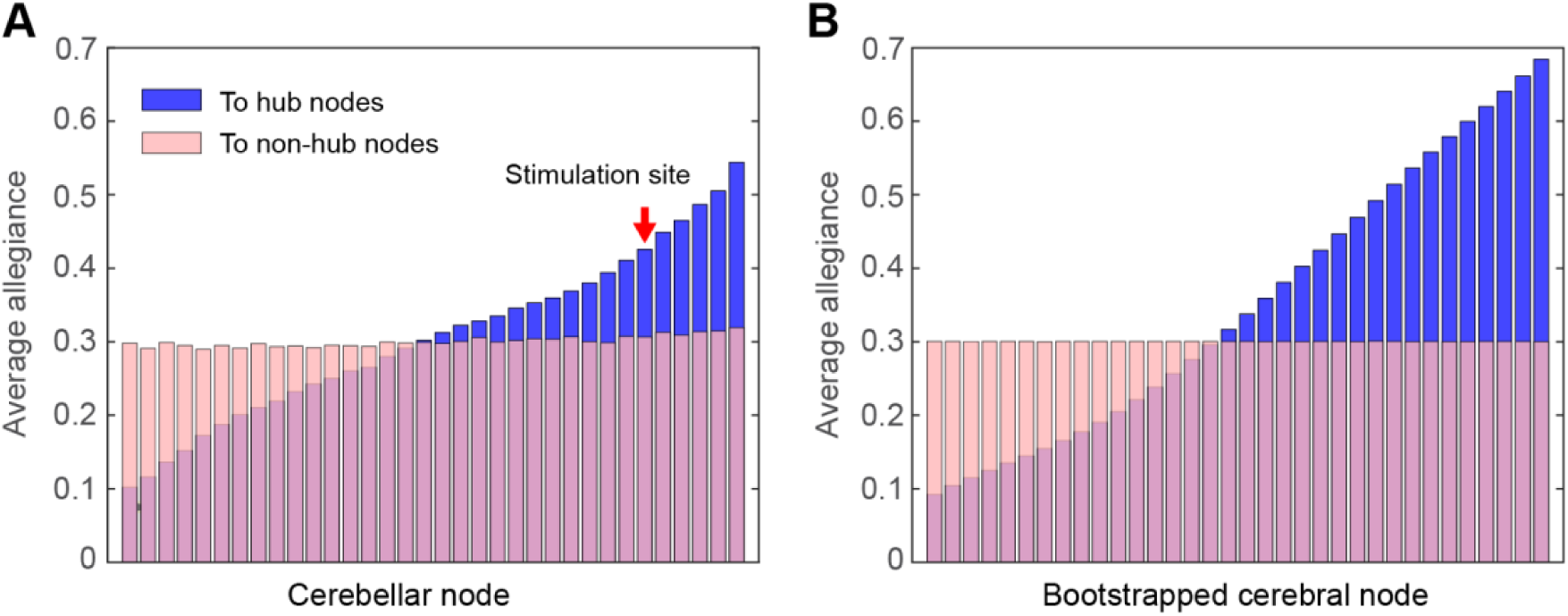
(A) Average allegiance of cerebellar nodes with hubs and non-hubs in the cerebral cortex. Hubs are defined as nodes with high overall allegiance (95%ile). Nodes are arranged in the order of increasing average allegiance to hub nodes. (B) Same as A, except that the equal number of nodes as cerebellum, i.e, 34, were picked within the cerebral cortex using a bootstrapping process 1000 times. Average allegiance values are computed by taking average across those bootstrapping iterations. Similar connectivity profiles for cerebellar and cerebral nodes are evident.

## Notes

### Competing Interest Statement

The authors have declared no competing interest.

